# Changes in Microbial Communities in Industrial Anaerobic Digestion of Dairy Manure Caused by *Caldicellulosiruptor* Pretreatment

**DOI:** 10.1101/2025.04.16.649167

**Authors:** Jakob Young, Maliea Nipko, Spencer Butterfield, Zachary Aanderud

**Affiliations:** Department of Biology, Brigham Young University, Provo, Utah 84602 USA; Department of Microbiology and Molecular Biology, Brigham Young University, Provo, Utah 84602 USA; Department of Sustainability & Renewable Energy Systems, University of Wisconsin-Platteville, Platteville, Wisconsin, 53818 USA; Department of Plant and Wildlife Sciences, Brigham Young University, Provo, Utah 84602 USA

**Keywords:** Co-occurrence network, community connectivity, biological pretreatment, syntrophic methanogenesis, acetoclastic methanogenesis, hydrogenotrophic methanogenesis

## Abstract

Hyperthermophilic pretreatment with *Caldicellulosiruptor* species (EBP) increases substrate availability in anaerobic digestion, but the effect on downstream microbial community composition in industrial systems is not characterized. Changes in microbial communities were determined at an industrial facility processing dairy manure in a modified split-stream system with three reactor types: 1) EBP tanks at 70–72°C, 2) mesophilic Continuously Stirred Tank Reactors (CSTRs), and 3) mesophilic Induced Bed Reactors (IBRs) receiving combined CSTR and EBP effluent. All reactors had a two-day hydraulic retention time. Samples were collected weekly for 60 days. pH, volatile fatty acid and bicarbonate concentrations, COD, and methane yield were measured to assess tank environmental conditions. Microbial community compositions were obtained via 16S rRNA gene sequencing. EBP pretreatment increased acetate availability but led to a decline in the relative abundance of acetoclastic *Methanosarcina* species in downstream IBRs. Rather, syntrophic methanogens, e.g., *Methanobrevibacter* species, increased in relative abundance and became central to microbial co-occurrence networks, particularly in association with hydrogen-producing bacteria. Network analysis also demonstrated that these syntrophic relationships were tightly coordinated in pretreated digestate but absent in the untreated CSTRs. By promoting syntrophic methanogenesis while increasing acetate concentrations, EBP pretreatment requires system configurations that enable acetoclast retention to prevent acetate underutilization and maximize methane yields.

**Importance:** Hyperthermophilic *Caldicellulosiruptor* pretreatment in anaerobic digestion increased acetate availability while suppressing acetoclastic methanogens and promoting robust syntrophic methanogenic networks. The results offer engineers microbial guidance essential for designing effective multi-stage industrial anaerobic digestion facilities employing EBP methods.

## 1. Introduction

Anaerobic digestion (AD) of dairy manure relies on microbial metabolism to synthesize biogas consisting almost entirely of CO_2_(g) and CH_4_(g). The conversion of organic waste into biogas proceeds through four sequential microbial groups: 1) hydrolytic bacteria, 2) acidogenic bacteria, 3) acetogenic and syntrophic bacteria, and 4) methanogenic archaea. The final step, methanogenesis, is performed by methanogenic archaea via three metabolic pathways. Primarily, acetoclastic methanogens, such as members of *Methanosarcina* and *Methanosaeta*, directly convert acetate into methane and bicarbonate in a single step catabolic reaction [1,2]:

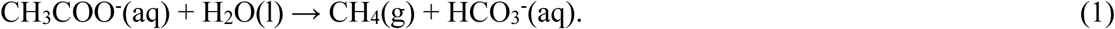

Alternatively, hydrogenotrophic methanogenesis involves two distinct steps, reliant on a syntrophic partnership between hydrogenogenic bacteria and hydrogenotrophic methanogens [1]. The bacteria initially oxidize acetate to produce hydrogen:

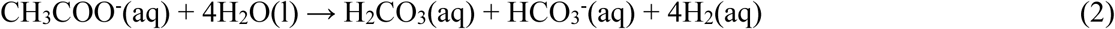

Then, via direct interspecies hydrogen transfer, hydrogenotrophic methanogens consume the hydrogen and produce methane [1,3–5]:

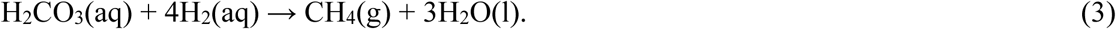

As Reactions 2 and 3 sum to Reaction 1, and each produces bicarbonate, the net effect of methanogenic activity is an increase in pH. Hydrogenotrophic pathways can also involve conversion of volatile fatty acids (VFAs) other than acetate. For example, butyrate is oxidized by acetogenic bacteria according to [6]:

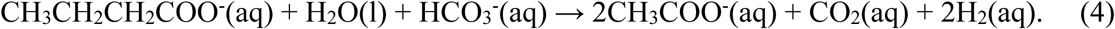

CO_2_ and H_2_ are then converted into methane by hydrogenotrophic methanogens via Reaction 3. Note that accumulation of acetate in digesters inhibits Reaction 4 by product inhibition. To avoid inhibition, substantial activity of acetoclastic methanogens is necessary.

Finally, methylotrophic methanogens degrade methylated compounds, such as methanol, to produce methane [7,8]. Similar to hydrogenotrophic methanogens, many methylotrophic methanogens require H_2_ for methanogenic activity, making them dependent on syntrophic interactions with bacteria [9]. Some methanogenic taxa, notably *Methanosarcina* spp., are metabolically flexible and can utilize a combination of acetoclastic, hydrogenotrophic, and methylotrophic methanogenesis depending on metabolite concentrations and environmental conditions [10].

Digester operation is typically monitored by total VFA concentration, pH, and alkalinity, all of which affect microbial activity and metabolic pathways. A pH range of 6.8-7.5 is optimal for methanogenesis [11,12]. Note that VFAs produced by hydrolysis and acidogenesis all exist as anions at pH 6.8-7.5. Dairy manure is buffered near pH 7.8 by ammonium bicarbonate, so it is critical to also monitor bicarbonate concentration. However, alkalinity as measured by titration with HCl includes both bicarbonate and volatile fatty acid anions (VFAAs). Total VFA concentration is commonly reported as acetate equivalents. For an accurate assessment of buffering capacity, the VFAA concentration must be subtracted from the measured alkalinity to estimate bicarbonate concentration.

Without pretreatment, biogas yields from AD are limited by incomplete hydrolysis of recalcitrant materials such as lignocellulose and peptidoglycans [13–15]. A variety of pretreatment methods have been explored to break down recalcitrant materials before digestion, such as using thermal conditions that accelerate rates of hydrolysis [16]. Specifically, the use of the extremophilic biological process (EBP) with *Caldicellulosiruptor* species has been shown to increase the rates of hydrolysis and acidogenesis, leading to greater biogas yields [6,17,18]. *Caldicellulosiruptor* species use exozymes to catalyze hydrolysis of lignin, celluloses, and peptidoglycans [19–21]. The saccharides released from celluloses are primarily catabolized into acetate. For example:

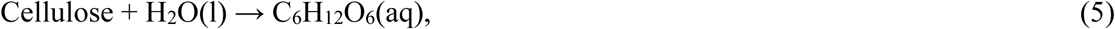

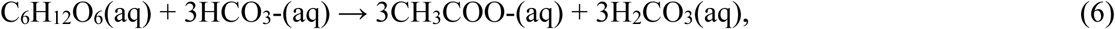

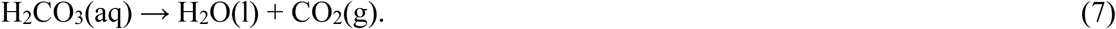

Studies have shown that pretreatment methods reshape AD microbial communities by enhancing hydrolysis and altering the chemical characteristics of digestate [22]. Specifically, thermal hydrolysis pretreatment often reduces overall microbial diversity, enriches ammonia-tolerant hydrogenotrophic methanogens, and strengthens syntrophic interactions during the anaerobic digestion of protein-rich substrates [23,24]. Consequently, hydrogenotrophic methanogenesis typically becomes the dominant form of methane generation under extreme pretreatment conditions, although pretreated reactors continue to produce methane through both acetoclastic and hydrogenotrophic pathways [22]. Metagenomic analyses of thermally pretreated sludge reactors demonstrated that higher temperatures correspond with a decrease in alpha diversity, while microbial networks become more interconnected [23]. However, most research has been confined to bench scale or single step pretreatment studies, leaving the longitudinal evolution of microbial communities and network connectivity in full scale multi-reactor digestion systems largely unexplored. As a result, plant engineers lack the microbial insights needed to optimize retention times, reactor configurations, and pretreatment strategies at industrial scale.

To address these limitations, we monitored microbial and chemical parameters at a commercial digestion facility using a modified split stream EBP pretreatment design. Our research group monitored the system but was not involved in the engineering design. At the facility, dairy manure was treated in EBP tanks inoculated with *Caldicellulosiruptor* species and maintained at elevated temperatures (70–72°C) to promote hydrolysis and acidogenesis. In parallel, mesophilic Continuously Stirred Tank Reactors (CSTRs) were fed raw manure without pretreatment. Effluents from both streams were combined and subsequently processed in mesophilic Induced Bed Reactors (IBRs) [25], combining acetate-rich digestate from pretreatment with an active acetoclastic community of methanogens from untreated digestate in the CSTRs. The hydraulic retention time (HRT) in all tanks was two days, as a means of maximizing throughput and maintaining continuous substrate availability. However, the low HRT likely impacted microbial retention by washout, especially of slow-growing methanogens. We expect the microbial relative abundance (RA) to represent a steady-state system determined by HRT, not abundances determined by growth limitations [26,27].

Given the combination of elevated temperatures and enhanced enzymatic hydrolysis provided by EBP tanks inoculated with *Caldicellulosiruptor* spp., we hypothesized four primary outcomes: (1) EBP pretreatment would increase lignocellulose hydrolysis rates, elevating acetate concentration in the digestate going into the IBRs; (2) The extreme thermal conditions in EBP tanks would force a microbial community shift, promoting thermotolerant methanogens, such as *Methanothermobacter* spp. and high temperature-tolerant bacterial populations; (3) Microbes that form biofilms would be favored due to the retention of particulate matter by the design of IBRs; and (4) The abundance of free-living acetoclastic methanogens would be depleted due to washout.

## 2. Materials and Methods

### 2.1 Site description

The study was conducted at a commercial facility with nine 189 cubic meter anaerobic reactors. Dairy manure was collected in a reception pit (RP) and fed into four extremophilic bacterial process (EBP) tanks at 70–72⁰C and two continuously stirred tank reactors (CSTRs) at 35–37⁰C. The effluent from all six initial tanks was mixed and fed into three induced bed reactors (IBRs) at 35–37°C. Each reactor had real-time monitoring of temperature, total biogas, and biogas methane content on site. Total methane production was calculated as the product of total biogas and percent methane. The EBP tanks were inoculated with a mixture of *C. bescii*, *C. owensensis*, *C. acetigenus*, and *C. saccharolyticus*. Hydraulic retention time in the EBP tanks, CSTRs, and IBRs was approximately two days.

### 2.2 Cultivation of Caldicellulosiruptor spp. and Inoculum Preparation

All four *Caldicellulosiruptor* spp. were acquired as lyophilized pellets from the German Collection of Microorganisms and Cell Cultures GmbH (DSMZ). Each culture was revived anaerobically following DSMZ guidelines until the colony reached approximately 2.5×10^6^ cells/mL. Specifically, *C. bescii* (DSMZ #6725) was grown in Medium 516 (DSMZ_Medium516.pdf), while *C. owensensis* (DSMZ #13100), *C. saccharolyticus* (DSMZ #8903), and *C. acetigenus* (DSMZ #12137) were each cultivated on Medium 640 (DSMZ_Medium640.pdf). Freezer stocks of each culture were prepared by mixing 500 µL of cell suspension (≈2.5 × 10^6^ cells/mL) with 500 µL of 60% glycerol in a Don Whitley Scientific miniMACS anaerobic chamber (West Yorkshire, UK). The resulting mixtures were gently agitated at room temperature for 10 minutes, allowing the glycerol to permeate the cells, and then stored at –80°C. For inoculum production, 2 mL each of *C. owensensis*, *C. acetigenus*, *C. saccharolyticus*, and *C. bescii* were added to 1 L of Medium 640 under anaerobic conditions in a miniMACS workstation, followed by incubation at 75°C until the population reached ≈2.5×10^6^ cells/mL. Cell density for all cultures was determined by microscopy. Each EBP tank received 2 L of inoculum for initial seeding.

### 2.3 Sample collection and storage

40 milliliters of sample were gathered from the effluent of each tank approximately weekly over 60 days. To ensure representative sampling, the reactor sample port was flushed prior to collection. Samples were frozen and transported in insulated packaging to the lab at BYU in Provo, Utah for chemical and biological analyses. Samples were kept frozen at −20⁰C until analysis. Note that this processing of samples likely increased the measured pH by removing CO_2_ from the solution.

### 2.4 Analyses of samples

A Hannah HI98195 probe (Hannah Instruments, Woonsocket, RI, USA) was used to measure pH. VFAs and chemical oxygen demand (COD) were measured with Hach TNT 872 Vial Tests and Hach TNT 822 Vial Tests, respectively (Hach, Loveland, CO, USA). The Hach TNT Vial Test for VFA measures total VFA and reports the result as acetate equivalents. For both VFA and COD measurements, each sample was subsampled, centrifuged for 10 minutes at 4000 RPM, and filtered with an Avantor VWR Syringe Filter (Glass fiber, 13mm, 1.0 um) (Avantor, Radnor Township, PA, USA). Because VFA concentrations in AD effluent tend to be too large for measurement by the Hach TNT Vial Test, each subsample was diluted 10-fold with Milli-Q ultrafiltered water prior to analysis.

Alkalinity was quantified via titration to pH 4 using 1 M HCl on a Hach Instruments TitraLab AT1000 (Loveland, CO, USA). Samples were diluted 100-fold with Milli-Q ultrapure water before titration. Because the measured alkalinity includes both bicarbonate and VFAAs, alkalinity was converted to an estimate of bicarbonate concentration (mg/L) with the equation:

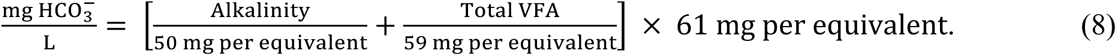

Alkalinity is expressed as mg CaCO₃ equivalents per liter and total VFA is expressed as mg acetate per liter. The constants 50 and 59 represent the mg of CaCO₃ and acetate that neutralize one mmole of H^+^ ions. The factor 61 is the molar mass of bicarbonate and is used to convert mmol to mg.

### 2.5 DNA extraction and sequencing

DNA was isolated from 300 mg of each frozen sample using the Qiagen DNEasy PowerWater Kit Pro (Qiagen, Kilden, Germany). Purity and concentration of the extracted DNA were checked with a Nanodrop One Microvolume UV-Vis Spectrophotometer (Thermo Fisher Scientific, Waltham, MA, USA). The 16S rRNA regions coded by the DNA were amplified for sequencing on an Illumina MiSeq platform (Illumina, San Diego, CA, USA), conducted at the Utah State University Sequencing Center in Logan, UT (https://cores.utah.edu/dna-sequencing/). The V4 hypervariable region of the 16S rRNA gene was targeted using primers 515FB and 806RB, producing ∼420 bp amplicons suitable for taxonomic profiling of bacterial and archaeal communities. First-round PCR reactions were performed in triplicate using Platinum II Hot Start master mix, followed by pooling and 1:50 dilution for indexing (ThermoFisher Scientific, Waltham, MA, USA). A second PCR added unique dual indices (p5/p7) to each sample (Integrated DNA Technologies, Coralville, IA, USA), and products were purified using SeraPure magnetic beads (Cytiva, Marlborough, MA, USA). Amplicon size was verified (∼420 bp) using a TapeStation or Fragment Analyzer (Agilent Technologies, Santa Clara, CA, USA), and samples were pooled stoichiometrically and sequenced on an Illumina platform (Illumina, San Diego, CA, USA).

### 2.6 DNA sequence processing

All 16S sequences were analyzed using the Qiime2 2022.2 pipeline [28]. First, raw reads were demultiplexed and subjected to quality filtering, removing chimeric reads and applying default denoising parameters via the DADA2 pipeline [29]. To standardize across samples, data were rarefied to 80,000 reads per sample. We built phylogenetic trees of denoised ASVs using fasttree2 [30]. ASVs were grouped at 97% similarity and assigned taxonomy via local BLAST against the SILVA taxonomic database [28,31]. Note that some species will not be detected or identified by the SILVA database [32]. Additionally, note that DNA presence in a tank does not denote activity [33]. All analyses were conducted at the ASV level and then aggregated to the genus or phylum level for visualization and interpretation.

For community dissimilarity, Bray-Curtis distances (Bray & Curtis, 1957) were calculated and visualized with Principal Coordinate Analysis (PCoA). Using the Qiime2R functionality within the tidyverse package, PCoA plots were generated in R v4.2.0 [34]. Permutational multivariate analysis of variance (PERMANOVA) using the adonis2 function in the vegan package further assessed the statistical significance of between-group differences [35].

### 2.7 Microbial correlation testing

Spearman correlations of all taxa abundance and environmental variables (COD, VFA, bicarbonate, pH, total methane production, and tank temperature) were determined. Spearman correlation was used because the microbial abundance data was non-normally distributed, often skewed, and compositional. Specifically, for each microbial taxon, abundance of the taxon was tested against COD, VFA, bicarbonate, pH, total methane production, and tank temperature across all samples from all tanks throughout the entire 60-day experiment. This allowed a comprehensive approach to correlation analysis, calculating correlation across the entirety of our data, rather than constraining the correlation to tanks that had changing environmental variables over time. In order to minimize Type I errors given the large number of correlations being tested, the p-values for these tests were adjusted using the Benjamini-Hochberg procedure.

### 2.8 Microbial co-occurrence networks

The microbial co-occurrence networks were calculated using the cooccur package in R [36]. All detected co-occurrences were filtered for significance (*p* < .05). Taxa unidentified by the SILVA database were removed from the analysis. Networks were calculated using the Fruchterman-Reingold method and a minimum distance was set between nodes to prevent overlap [37]. Distance between nodes was set to the inverse of effect size and the size of nodes was set to node connectivity. The calculated network was plotted using ggraph on R [38]. Connectivity, eigenvector values, and betweenness were calculated for each node. Modularity and connectivity statistics were computed for each graph.

## 3. Results and Discussion

### 3.1 Community compositions

#### 3.1.1 Reception Pit (RP)

The RP tank accepts raw dairy manure that has a stable microbial community (Figure 1, Figure 2, Table 1, Table 2). Hydrogenotrophic *Methanobrevibacter* (63.7±2.3%) and members of Methanobacteriaceae (33.5±2.0%) together dominated the methanogen community. *Methanosarcina* genus (0.3±0.2%), capable of acetoclastic, methylotrophic, and hydrogenotrophic methanogenesis, was relatively low in abundance. Prior comparative metagenomic analysis observed that hydrogenotrophic archaea typically dominate the archaeal population within raw manure, while acetoclastic populations remain minor [39]. *Methanosphaera* comprised an average RA of 2.4% of the methanogenic community in the raw manure (Table 2).

**Figure 1.**
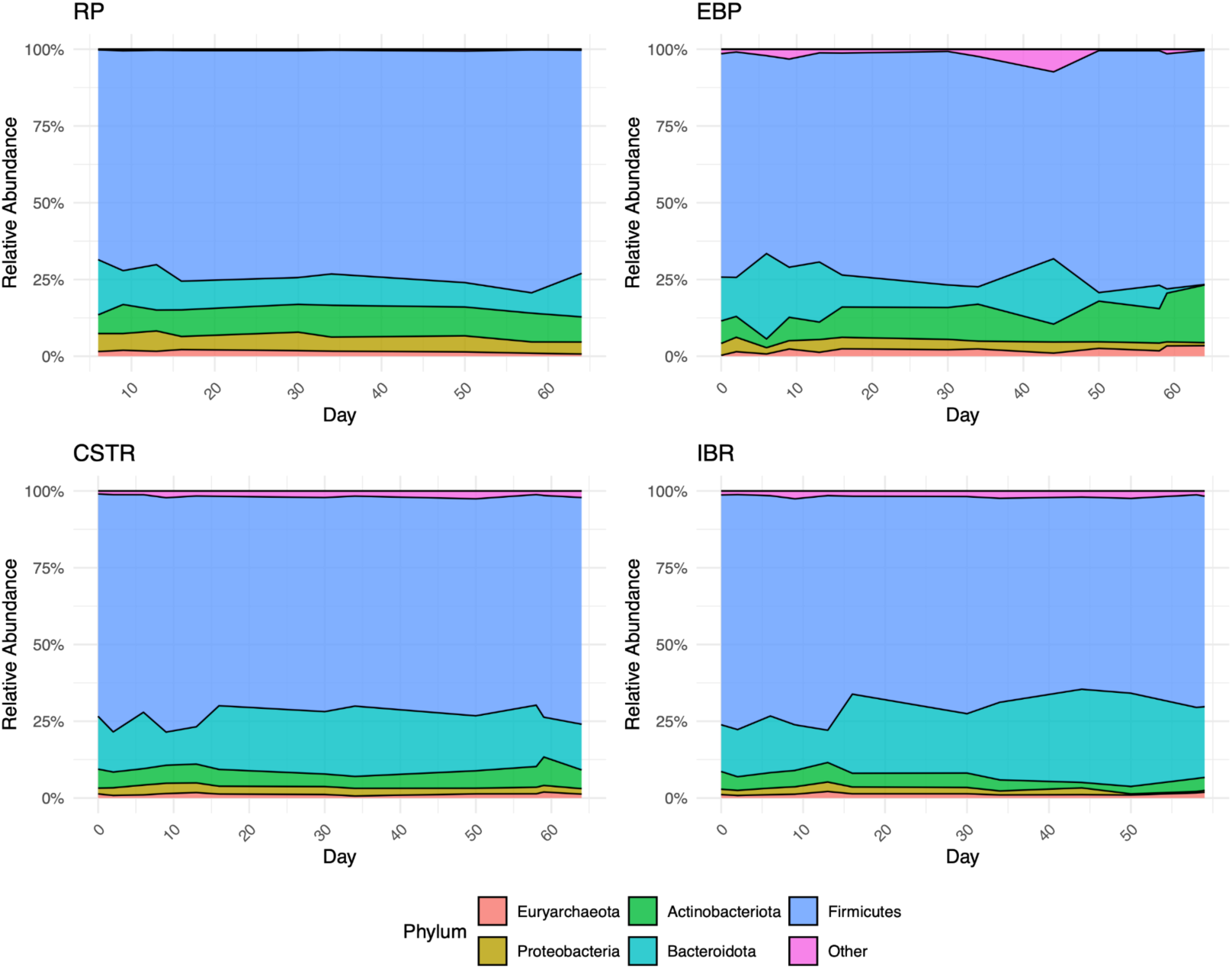
Stacked area plots showing relative abundance of the five most common microbial phyla in each tank type over time. The abundance values represent the average across all tanks within each category by date. See Table 1. Bacteroidota relative abundance decreases in the EBP tanks and increases in the IBRs. Actinobacteria relative abundance increases in the EBP tanks.

**Figure 2.**
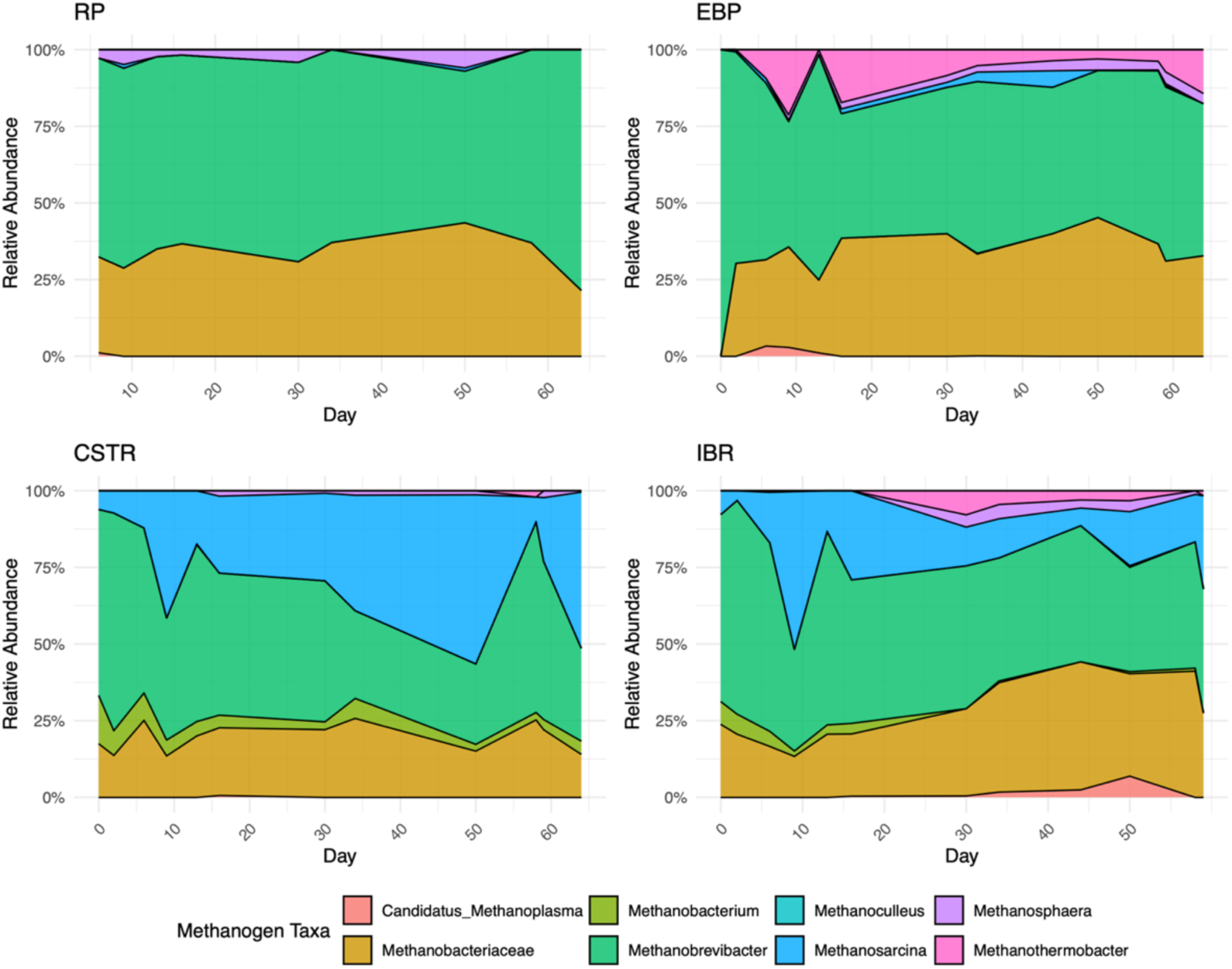
Stacked area plot depicting the community abundance of methanogenic archaea over time across the three tank categories and the Reception Pit (RP). Abundances are averages across all tanks within the category on that date. See Table 2. Hydrogenotrophic methanogens dominate all communities. Acetoclastic Methanosarcina relative abundance grows in the CSTRs and declines in the IBRs. Methylotrophic and thermophilic methanogen relative abundance grows in the IBRs.

**Table 1.** Summary of non-methanogenic community composition by mean relative abundance of phyla across samples, separated by RP (n = 10), EBP tanks IBRs (n = 33), and CSTRs (n = 23). Since the relative abundance is averaged from all samples throughout the experiment, standard error is included. See Figure 1.

| Phylum | RP<br>1 tank, n =<br>10 | EBP<br>4 tanks, n =<br>37 | CSTR<br>2 tanks, n =<br>23 | IBR<br>3 tanks, n =<br>33 |
| --- | --- | --- | --- | --- |
| Firmicutes | 66.8% ± 6.5 | 72.9% ± 1.1 | 72.0% ± 0.8 | 70.3% ± 0.9 |
| Bacteroidota | 11.6% ± 1.2 | 10.4% ± 1.6 | 16.9% ± 1.0 | 20.8% ± 1.2 |
| Actinobacteriota | 9.7% ± 1.2 | 10.5% ± 0.9 | 5.8% ± 0.3 | 4.3% ± 0.3 |
| Proteobacteria | 9.6% ± 4.6 | 2.6% ± 0.2 | 2.5% ± 0.1 | 1.8% ± 0.2 |
| Euryarchaeota | 1.4% ± 0.2 | 2.2% ± 0.2 | 1.3% ± 0.1 | 1.3% ± 0.1 |
| Other | 0.8% ± 0.4 | 1.5% ± 0.4 | 1.5% ± 0.1 | 1.5% ± 0.1 |

**Table 2.** Summary of methanogen community composition by mean relative abundance across samples, separated by tank type. The relative abundance is presented as the percentage of the methanogenic community. Since the relative abundance is averaged from all samples throughout the experiment, standard error is included. See Figure 2.

| **Methanogen** | **RP**<br>1 tank, n = 10 | **EBP**<br>4 tanks, n = 37 | **CSTR**<br>2 tanks, n = 23 | **IBR**<br>3 tanks, n = 33 |
| --- | --- | --- | --- | --- |
| *Methanobrevibacter* spp. | 63.7% ± 2.3 | 56.3% ± 2.7 | 47.8% ± 3.5 | 48.7% ± 2.35 |
| Members of Methanobacteriaceae | 33.5% ± 2.0 | 33.5% ± 1.7 | 19.9% ± 1.2 | 26.5% ± 1.74 |
| *Methanosarcina* spp. | 0.3% ± 0.2 | 1.2% ± 0.4 | 25.6% ± 4.0 | 16.8% ± 2.49 |
| *Methanothermobacter* spp. | ND<sup>1</sup> | 6.2% ± 1.5 | 0.1% ± 0.1 | 1.6% ± 0.57 |
| *Methanobacterium* spp. | ND | ND | 5.5% ± 0.9 | 2.4% ± 0.5 |
| *Methanosphaera* spp. | 2.4% ± 0.7 | 2.0% ± 0.3 | 0.5% ± 0.2 | 1.4% ± 0.4 |
| Members of Candidatus *Methanoplasma* | 0.1% ± 0.1 | 0.4% ± 0.2 | ND | 1.0% ± 0.4 |

<sup>1</sup> ND = Not Detected

#### 3.1.2 Continuously Stirred Tank Reactors (CSTRs)

The CSTRs, fed manure from the RP, exhibited stable microbial communities of Firmicutes (72.0±0.8%), Bacteroidota (20.8±1.2%), and Actinobacteriota (4.3±0.3%). The main methanogenic taxa were *Methanobrevibacter* (47.8±3.5%), *Methanosarcina* (25.6±4.0%), and members of Methanobacteriaceae (19.9±1.2%). In the mesophilic CSTRs, the RA of *Methanosarcina* genus grew relative to the hydrogenotrophic methanogens dominant in the raw manure. Members of *Methanosarcina* exhibit faster acetate consumption kinetics and higher tolerance for elevated acetate concentrations, allowing them to outcompete hydrogenotrophic methanogens under acetate-rich conditions [40,41]. Additionally, over time, the relative abundance of *Methanosarcina* further increased in the CSTRs (Figure 2). *Methanosarcina* was correlated with lower VFA (Spearman’s ρ = −0.35, p < .01, n = 103) and higher bicarbonate (Spearman’s ρ = 0.55, p < .001, n = 103) concentrations. In the CSTRs, therefore, *Methanosarcina* spp. actively facilitated acetoclastic methanogenesis, consuming VFAs and releasing bicarbonate, consistent with Reaction 1.

#### 3.1.3 Extremophilic Biological Process (EBP)

Firmicutes abundance was constant, comprising the majority of microbes in these tanks (72.9±1.1%), Figure 1. The presence of the Thermotogota and Fusobacteriota phyla, which fall under the other category in Figure 1, were noted on occasion. Fusobacteriota specializes in degradation of plant biomass such as cellulose and cellobiose and likely contributed to the hydrolysis and metabolism of recalcitrant compounds in the EBP tanks [42]. Thermotogota are known to prefer higher temperatures and produce H_2_, possibly forming relationships with syntrophic methanogens [43,44]. In this case *Methanosarcina* was undetected in the EBP tanks, likely because the growth of these archaea are inhibited above 65°C [45,46]. As noted above, the split-stream configuration delivers CSTR effluent into the IBRs precisely to replenish the mesophilic acetoclastic methanogens inactivated by the hyperthermophilic EBP pretreatment. As seen in Table 1, the hyperthermophilic EBP reactors have a larger average RA of Euryarchaeota (2.2±0.2% of total) compared with the CSTRs and IBRs. This is explained by two sources of methanogen DNA. Microorganisms from RP have died or gone dormant and their genetic material is detected [47]. Additionally, thermophilic methanogens, e.g. *Methanothermobacter* spp. (6.2±1.5% of methanogens in EBP), from the raw manure grew in the EBP tanks. The growth in RA of *Methanothermobacter* genus and members of Thermotogota phylum confirms our hypothesis that hyperthermophilic pretreatment selects for thermotolerant methanogens and bacteria. Note that *Caldicellulosiruptor* spp. are present but not included in the analysis.

#### 3.1.4 Induced Bed Reactors (IBR)

In the IBR tanks fed a mixture of effluents from the CSTRs and EBP tanks, the RA of Bacteroidota (20.8±1.2%) increased over time, seemingly at the expense of Firmicutes (70.3±0.9%) and Actinobacteriota (4.3±0.3%) (Figure 1, Table 1). The dominant methanogenic taxa in the IBRs were *Methanobrevibacter* (48.7±2.4%), *Methanosarcina* (16.8±2.5%), and members of Methanobacteriaceae (26.5±1.7%). The RA of *Methanosarcina* in the IBRs was smaller than that of the CSTRs and declined over time (Figure 2). This decline confirms our hypothesis that many members of *Methanosarcina*, free-living acetoclasts, would be depleted under low HRT due to microbial washout. In contrast, the RA of both methylotrophic methanogens (e.g., Candidatus *Methanoplasma,* 1.0±0.4%), and hydrogenotrophic methanogens (e.g., *Methanothermobacter*, 1.6±0.6%), from the raw manure grew in the IBRs. These findings denote a decline in the acetoclastic pathway relative to alternative methanogenesis pathways. Additionally, the RA of both Candidatus *Methanoplasma* (Spearman’s ρ = 0.48, p < .001, n = 54) and *Methanothermobacter* (Spearman’s ρ **=** 0.48, *p* < .001, n = 54) correlated with higher methane production. The evidence from the IBRs indicates EBP pretreatment shifts the methanogenic community away from acetoclastic methanogenesis, likely because IBRs retain large solid particles but allow soluble substrates, such as acetate, to pass through underutilized due to low HRT.

### 3.2 Co-occurrence networks

#### 3.2.1 Reception pit (RP)

The RP co-occurrence network was distinct due to its high modularity of 0.74 (Table 3). The community is loosely organized and contains many disconnected groups of taxa (Figure 3). Methylotrophic *Methanosphaera* and hydrogenotrophic *Methanobrevibacter* are noted in the manure community network but are relatively disconnected and possess low eigenvector values. The average taxon had a positive relationship with 2.7 other taxa.

**Table 3.**
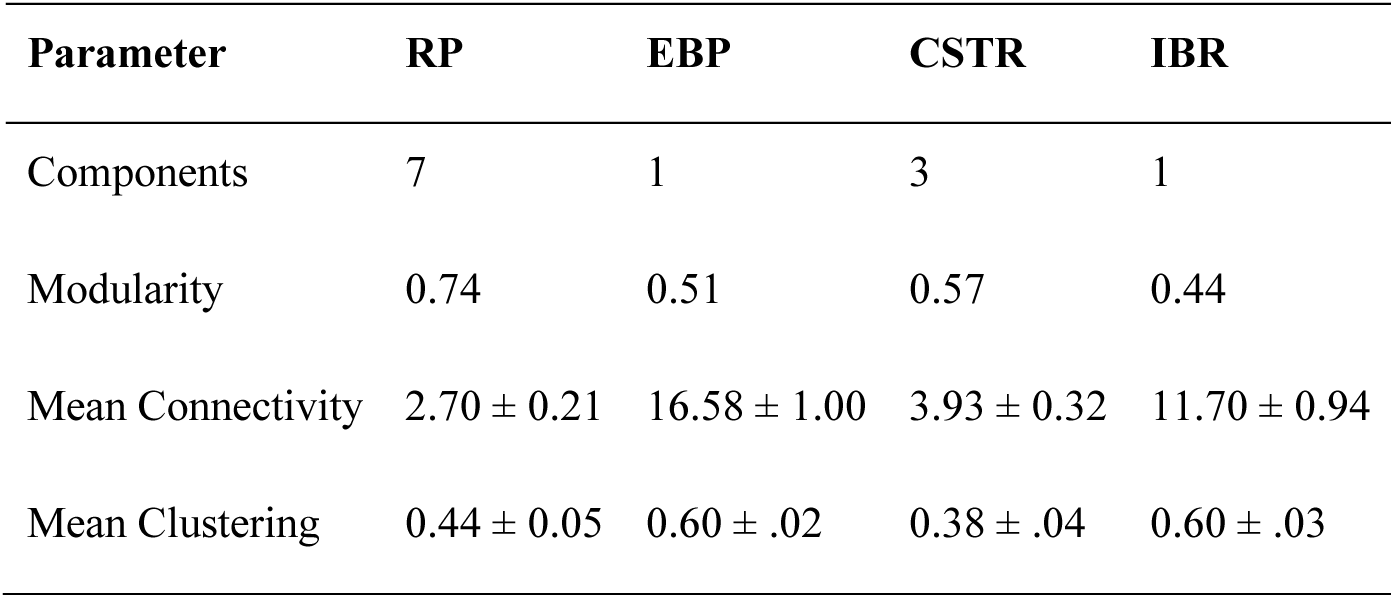
Co-occurrence network summary statistics by tank type. Components refer to the number of disconnected groups of taxa in a community. Modularity measures how well a network divides into communities with dense internal and sparse external connections. Mean Connectivity is the average number of taxa each taxon is connected to within a network. Mean clustering represents the degree to which nodes in a network tend to cluster together into tight groups. Means are presented ± SE.

| Parameter | RP | EBP | CSTR | IBR |
| --- | --- | --- | --- | --- |
| Components | 7 | 1 | 3 | 1 |
| Modularity | 0.74 | 0.51 | 0.57 | 0.44 |
| Mean Connectivity | $2.70 \pm 0.21$ | $16.58 \pm 1.00$ | $3.93 \pm 0.32$ | $11.70 \pm 0.94$ |
| Mean Clustering | $0.44 \pm 0.05$ | $0.60 \pm .02$ | $0.38 \pm .04$ | $0.60 \pm .03$ |

**Figure 3.**
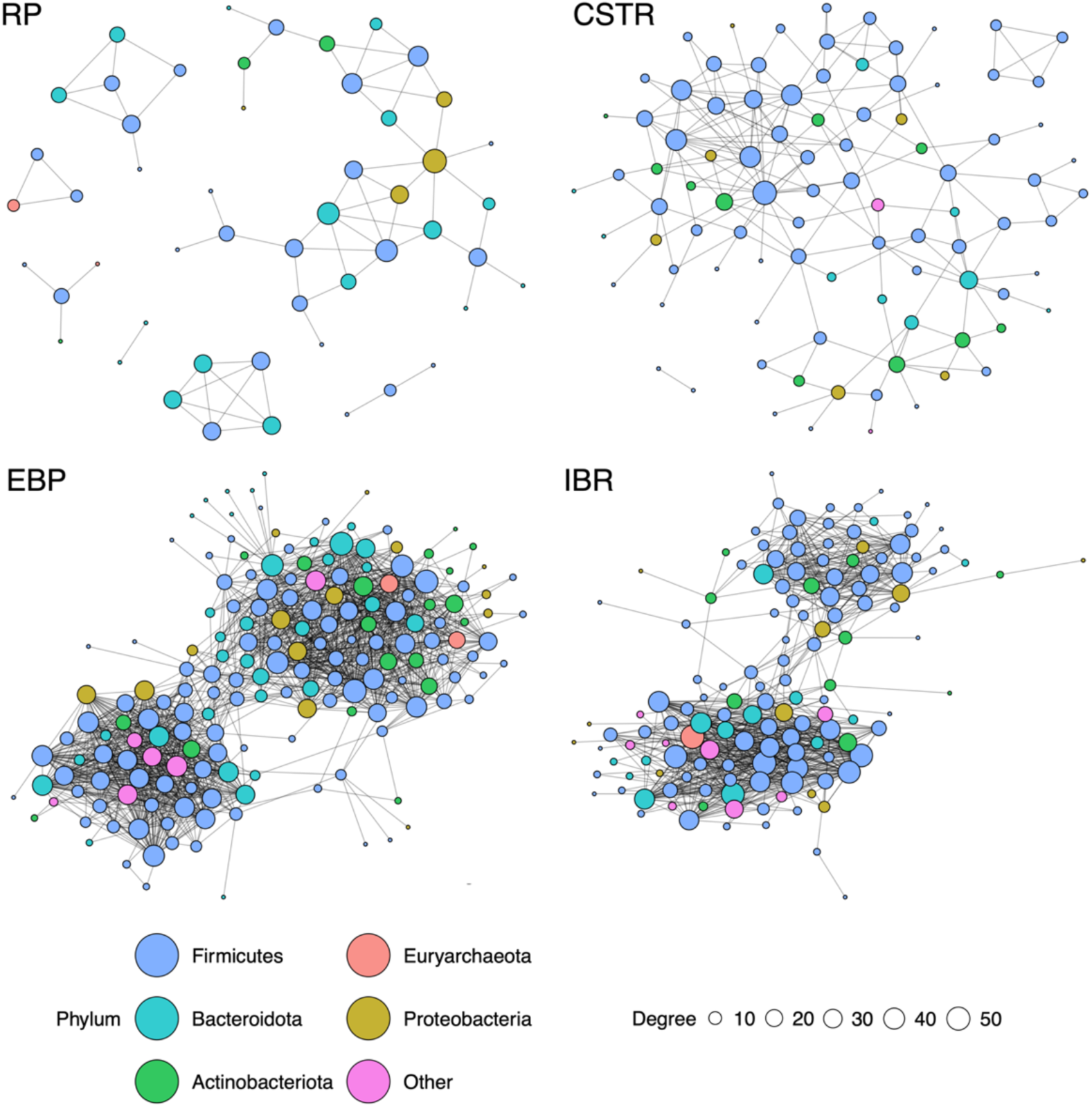
Co-occurrence network graphs depicting community relationships between microbial taxa in the RP (n = 10), EBP tanks (n = 37), IBRs (n = 33), and CSTRs (n = 23). Lines are drawn between taxa when they are observed together in samples significantly more frequently than random (p < .01). The closeness between a pair of taxa corresponds with the effect size of the relationship. The size of the node represents the number of connections to other taxa. See Table 3 for network statistics. EBP and IBR networks are dense and interconnected, with hydrogenotrophic methanogens central to their communities. Raw manure (RP) and CSTR communities are disconnected and indistinct. No co-occurrence relationships with methanogenic taxa were detected in the CSTR tanks.

#### 3.2.2 CSTRs

The CSTR network had a modularity value of 0.57 and consisted of 3 components (Table 3). The most influential taxa with the highest eigenvector values were entirely of the *Firmicutes* phylum. Despite having an average RA of 1.3% (Table 1) in the CSTR tanks, no positive relationships toward methanogenic taxa were detected. Many known acetogenic taxa, such as members of Lachnospiraceae and *Syntrophomonas*, presented with high betweenness factors [48,49]. Acetogenic taxa therefore played a key role in this community, transforming the products of related hydrolytic and acidogenic bacteria into acetate. The lack of hydrogenotrophic methanogens in the network suggests a fundamental limitation to methane production due to their reliance on interspecies hydrogen transfer [4].

#### 3.2.3 EBP tanks

The EBP pretreatment tank co-occurrence network exhibited a highly dense and interconnected community (Figure 3). Each taxon in the EBP tanks was connected to an average of 16.6 other taxa (Table 3). The network had a modularity value of 0.5. Members of Firmicutes and Bacteroidota, including *Herbinix* and *Petrimonas* genera, comprised the majority of taxa with high eigenvector values. Notably, the third most influential taxon in the EBP tanks was a member of the Verrucomicrobiota phylum, presenting with an eigenvector of 0.995. The role of Verrucomicrobiota is largely uncharacterized in the context of AD. Methylotrophic *Methanosphaera* and hydrogenotrophic *Methanobrevibacter* displayed centrality in the community, both exhibiting high eigenvector and degree values. Both *Methanosphaera* and *Methanobrevibacter* exhibited strong co-occurrence relationships with several syntrophic bacteria capable of H_2_ production, notably members of Thermoanaerobacteraceae, Thermovenabulum, and *Caldicoprobacter* [50–52].

In thermophilic reactors, enhanced breakdown of proteins can lead to increased free ammonia accumulation [53,54]. When ammonia levels rise under high-temperature conditions, acetoclastic pathways can be suppressed and digestion relies more heavily on syntrophic interspecies hydrogen transfer [55,56]. It is therefore unsurprising that, *Methanosarcina*, the only genus capable of acetoclastic methanogenesis detected in the system, was detected in very low abundance (Table 2) and did not display positive co-occurrence with any identified bacterial taxa. In addition to acetoclastic suppression, both thermophilic temperatures and short retention times in AD have been seen to cause a shift towards hydrogenotrophic methanogenesis [5,57]. In this case, increased free ammonia from hyperthermophilic conditions under EBP and the brief HRT hindered acetoclastic pathways and shifted the community towards syntrophic methanogenic networks.

#### 3.2.4 IBRs

In the IBRs processing the mixed CSTR and EBP effluents, the co-occurrence network exhibited a modularity of 0.43 and a mean connectivity of 11.69 taxa per node (Table 3). *Methanosphaera* displayed both the highest betweenness and a high eigenvector score (0.8), marking it as a keystone species that sustained a highly interconnected syntrophic web with diverse bacterial partners. In particular, due to its hydrogen dependency, *Methanosphaera* formed strong relationships with syntrophic bacteria known to produce H_2_, such as members of Thermoanaerobacteraceae, *Defluviitoga*, and *Caldicoprobacter* [50,51,58,59].

The persistence of these syntrophic taxa in the IBR network, originally structured under hyperthermophilic EBP pretreatment, highlights the resilience of the hydrogen-dependent community when conditions shifted to mesophilic temperatures. No methanogens capable of acetoclastic methanogenesis presented in the co-occurrence network. Despite elevated VFAA concentrations in the IBRs, no difference was discerned between the bicarbonate concentrations of the CSTRs and IBRs (Tukey, Δ = −198.53, p > 0.05). Under active acetoclastic conversion of the surplus acetate (Reaction 1), we would expect to see elevated bicarbonate levels in the IBRs. Acetoclastic pathways were therefore insufficient to convert the excess acetate into additional methane and bicarbonate, suggesting that elevated methane yields in these mesophilic reactors relied on the enduring syntrophic interactions formed during thermophilic pretreatment. The continued centrality of syntrophic taxa in the IBR networks, including *Methanosphaera*, supports our hypothesis that biofilm-forming microorganisms are favored under IBR retention conditions.

#### 3.2.5 Comparison of networks

The RP network was the most disconnected, exhibiting the highest modularity (0.74) and lowest mean connectivity value (2.7). Despite slightly higher connectivity, the CSTR network remained structurally similar to the RP. In contrast, the EBP and IBR communities exhibited much higher connectivity and resembled each other. The EBP tanks have the highest connectivity value (16.6) of all tanks. Both the EBP and IBR networks consisted of 1 component, while the RP and CSTR consisted of multiple components. The EBP and IBR networks have clustering coefficients of 0.6, larger than both the RP (0.44) and CSTRs (0.38). While the most influential members of the CSTR network were entirely Firmicutes, the most influential taxa in both the EBP and IBR networks were more diverse, consisting of Firmicutes, Bacteroidota, and others. The syntrophic webs assembled during hyperthermophilic pretreatment persisted into the IBRs, demonstrating the resilience of hydrogenotrophic partnerships even under mesophilic operation. These persistent networks remained central to community connectivity and function downstream. Notably, the IBR network had the lowest modularity value (0.43), indicating that its community was the least segmented into disconnected subgroups among all tank types. Except the mesophilic CSTRs processing raw manure, syntrophic methanogens exhibited relationships with other taxa in all other networks. Acetoclastic *Methanosarcina* spp. did not have co-occurrence relationships in any of the tanks due to the high levels of acetate from the RP and EBP pretreatment. Together, these patterns indicate that EBP pretreatment causes large changes in downstream microbial networks, favoring tightly connected syntrophic communities and diminishing the role of free-living acetoclastic methanogen species. From an operational standpoint, connectivity and community structure may be useful indicators of functional dominance, guiding adjustments to HRT in order to maximize methane production.

### 3.3 Community dissimilarity and relationship with environmental variables

Dissimilarities between microbial community compositions, i.e. all of the detected taxa in a sample weighted according to their RA, were assessed between tank types using PCoA (Figure 4). The strength and direction of the correlations among significant environmental variables and community compositions is overlaid as vectors. Microbial communities differed by tank type (PERMANOVA, R² = 0.36, p < .001) due to large differences in environmental conditions between tank types. Axis 1, with 40.42% variance explained, largely differentiated communities by VFA and bicarbonate concentrations. Additionally, axis 1 horizontally separated the reactors responsible for methane production (CSTRs and IBRs) from the raw manure and pretreatment tanks (RP and EBP tanks). Axis 2, with 13.29% variance explained, vertically differentiated communities by pH, COD, temperature, and TDS.

**Figure 4.**
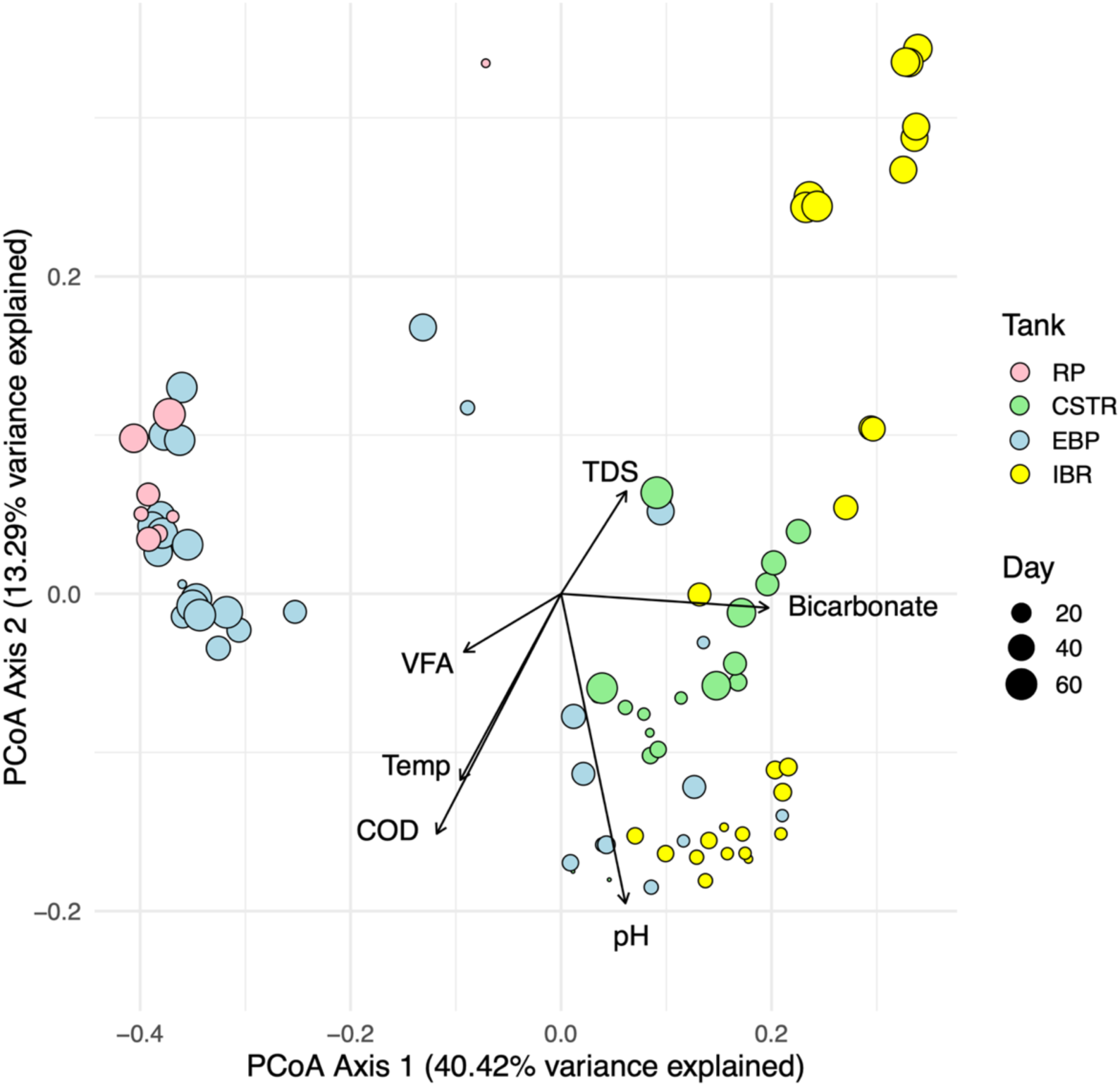
Principal Coordinates Analysis (PCoA) biplot of microbial community composition in all samples with colors corresponding to tank type. Point sizes indicate when the sample was collected with smaller points represent samples collected near the beginning of the experiment while the larger points represent samples collected near the end of the experiment. A point represents the microbial community composition of a sample, i.e. all of the detected taxa in a sample weighted according to their relative abundance. Vectors depict significant correlations with environmental variables and microbial community compositions. The length of each vector indicates the strength of the correlation with each of the environmental variables, total dissolved solids (TDS), volatile fatty acids (VFA), temperature (Temp), pH, and bicarbonate. Reactors responsible for methane production (CSTRs and IBRs) and tanks responsible for pretreatment and storing manure (EBP tanks and RP) are separated by Axis 1. Both the EBP and IBR communities experienced change over time, are correlated with decreasing pH. EBP tanks and IBRs were similar in the beginning but diverged with time. EBP tanks followed the diagonal from the lower right to upper left. IBRs followed a vertical path, aligned with axis 2, moving from the bottom right to the top right. RP and CSTR community compositions did not substantially change over time.

The trajectory of the EBP tanks over time aligns with expected *Caldicellulosiruptor* metabolic activity. Near the beginning of the experiment, communities in the EBP tanks were associated with higher COD, temperature, and pH. But these communities progressively became associated with high VFA and low bicarbonate concentrations as time passed, consistent with *Caldicellulosiruptor* metabolism outlined in Reactions 5, 6, and 7. This trend is further supported in the environmental data, with the EBP tanks maintaining the highest acetate and the lowest bicarbonate levels across all tank types (Figure 5), supporting our first hypothesis. During the experiment, the EBP communities evolved to become similar to the raw manure communities in the RP (Figure 4).

**Figure 5.**
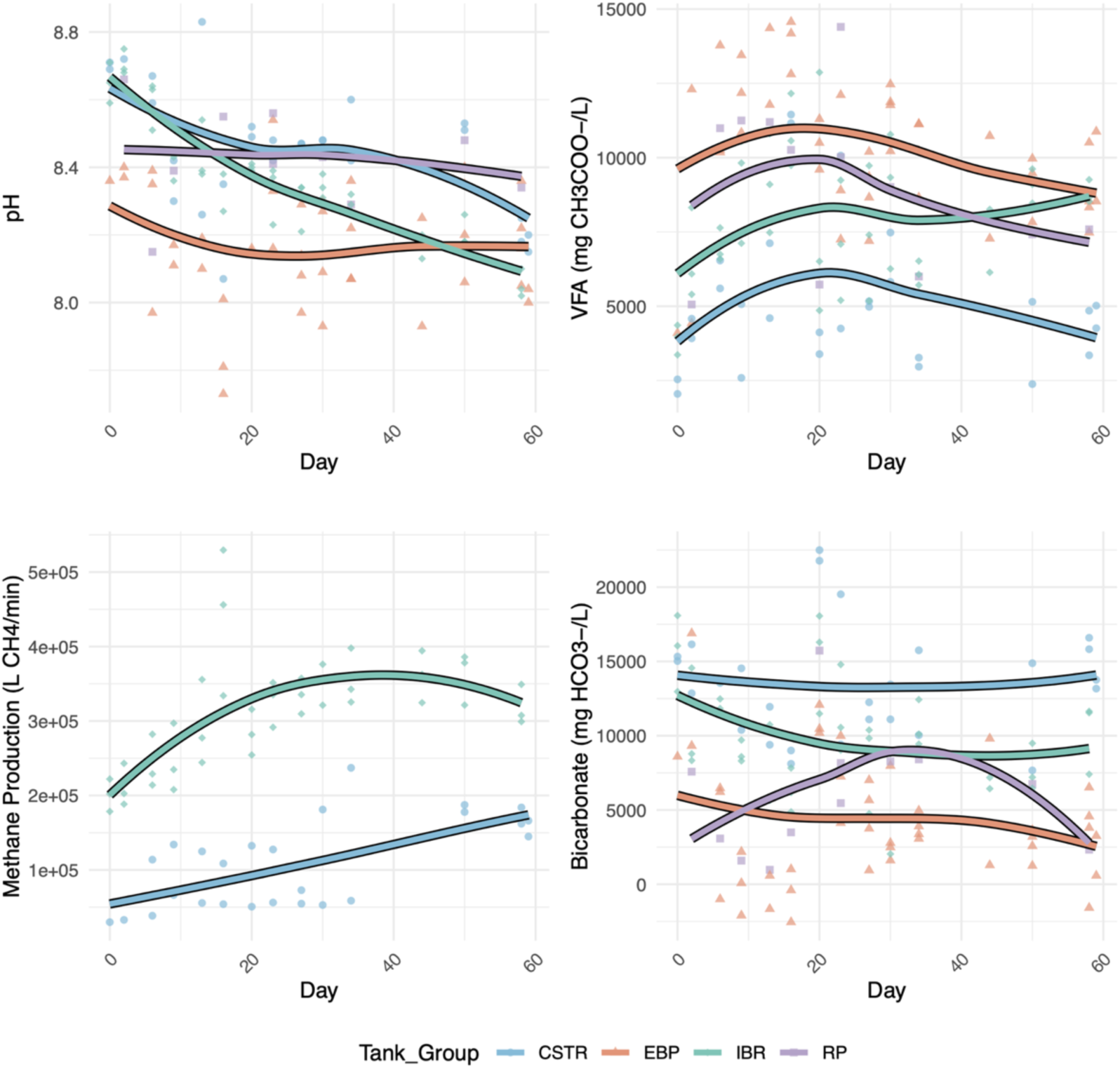
Trends in pH, Total VFA, bicarbonate, and methane gas flow (L/min) by tank type over time. Regressions are calculated and drawn using the Locally Estimated Scatterplot Smoothing (LOESS) technique to clearly illustrate trends in variables over time. EBP tanks consistently had lower pH and bicarbonate, and higher total VFAA concentrations than other tanks. IBRs exhibited decreasing pH and increasing VFAA concentrations over time. CSTRs had higher bicarbonate concentrations and lower total VFA concentrations than other tanks. IBRs consistently produced more methane on average than the CSTRs.

Likewise, movement of the CSTR and IBR tanks on the dissimilarity matrix demonstrates methanogenic activity. Specifically, CSTR and IBR communities were notably different from those of the RP and EBP and were distinctly correlated with higher bicarbonate concentrations, indicating active methanogenic communities, i.e., see Reactions 1, 2, and 3. The environmental data provides further evidence of this, with bicarbonate levels remaining higher in both the CSTR and IBR tanks than the RP and EBP tanks (Figure 5). At the start of the experiment, the CSTRs and IBRs were similar (Figure 4), but the IBR communities became distinct due to introduction of EBP effluent, dominated by thermophiles not suitable for activity in the mesophilic IBRs. Consequently, the microbial community that developed in the IBRs over time was shaped by both the physiochemical characteristics of the EBP effluent and the community in the CSTR effluent, ultimately resulting in a community distinct from both the EBP and CSTR communities. The divergence of IBR communities over time highlights how upstream pretreatment and reactor design directly shaped downstream function and community structure. Also, as the experiment progressed, the IBR communities became more correlated with increased TDS and decreased pH. The IBR tanks saw a unique increase in acetate and decrease in bicarbonate concentrations over the course of the experiment (Figure 5). These findings are explained by a large increase in syntrophic activity, which replaces bicarbonate with acetate as seen in Reaction 2. While the IBR tanks exhibited higher methane yields than the CSTRs, the large concentration of acetate received from EBP pretreatment remained critically underutilized (Figure 5). The brief HRT in the IBR tanks prevented the growth and accumulation of acetoclastic methanogens responsible for the conversion of acetate into bicarbonate which would stabilize pH (Reaction 1). Lastly, the design of IBRs retains large particles which accounts for the correlation with increased TDS.

## 5. Conclusions

EBP pretreatment caused acetate accumulation and elevated methane production in downstream IBRs with two-day HRT. The RA of *Methanosarcina*, the only genus capable of acetoclastic methanogenesis detected in the system, increased substantially in the CSTRs processing raw manure. However, the RA of *Methanosarcina* declined in the IBRs, despite being introduced continuously from the CSTRs alongside high levels of acetate from the EBP tanks. The continued decline of *Methanosarcina* highlights the importance of aligning pretreatment strategies with system design, such as HRT. In this case, the low HRT in the IBRs limited the persistence of acetoclastic methanogens and allowed acetate to pass through the reactors underutilized. These results suggest that the higher methane yield observed in the IBRs was driven not by acetoclasts, but by growth in both RA and connectivity of syntrophic methanogens, especially members of Methanobacteriaceae.

Co-occurrence network analysis proved to be a useful method of evaluating microbial functionality in complex AD systems. Compared to the RP and CSTR tanks, the EBP and IBR networks displayed considerably higher connectivity and clustering, reflecting more cohesive microbial communities centered on syntrophic methanogenic activity. Within both EBP and IBR networks, *Methanobrevibacter* and *Methanosphaera* acted as central taxa, forming strong associations with known H₂-producing bacteria including members of *Caldicoprobacter*. These interactions demonstrate how syntrophic methanogenic pathways can compensate for loss of acetoclastic activity under certain conditions. In contrast, the CSTR networks lacked associations between hydrogenotrophic methanogens and bacteria, suggesting functional constraints in systems relying on acetoclastic dominance.

Together, these results demonstrate that thermophilic pretreatment with *Caldicellulosiruptor* species substantially alters environmental parameters, such as elevating acetate concentrations, while simultaneously enhancing downstream syntrophic pathways of methanogenesis. System designers should account for these microbial community changes by pairing pretreatment with reactor configurations and retention times that support the growth and retention of slow-growing methanogenic taxa, especially acetoclasts. Additionally, network connectivity analysis can serve as a valuable indicator for monitoring microbial metabolic activity and shifts in methanogenic pathways in full-scale AD systems.

## Author Contributions

Conceptualization, Zachary Aanderud, Maliea Holden, and Jakob Young; methodology, Zachary Aanderud; validation, Zachary Aanderud and Maliea Holden; formal analysis, Jakob Young; investigation, Jakob Young and Zachary Aanderud; resources, Zachary Aanderud; data curation, Jakob Young and Spencer Butterfield; writing—original draft preparation, Jakob Young; writing— review and editing, Jakob Young, Maliea Holden, and Spencer Butterfield; visualization, Jakob Young; supervision, Zachary Aanderud; project administration, Zachary Aanderud; funding acquisition, Zachary Aanderud. All authors have read and agreed to the published version of the manuscript.

## Data Availability Statement

Data is available upon request.

## Acknowledgments

The authors thank Lee D. Hansen and Jaron C. Hansen for their assistance with both editing and experimental design. We also acknowledge the help of Jane Allen, Celeste Doxey, Katie Tano, Meredith Murdock, Kailey Condie, and Lauren Membreño with sample processing.

## Abbreviations

The following abbreviations are used in this manuscript:

RP: Reception Pit
CSTR: Continuously Stirred Tank Reactor
EBP: Extremophilic Biological Process
IBR: Induced Bed Reactor
AD: Anaerobic Digestion
VFA: Volatile Fatty Acid
VFAA: Volatile Fatty Acid Anion
COD: Chemical Oxygen Demand
TDS: Total Dissolved Solids
RA: Relative Abundance
HRT: Hydraulic Retention Time

